# Two Receptor-Like Kinases Required For Arabidopsis Endodermal Root Organisation Shape The Rhizosphere Microbiome

**DOI:** 10.1101/816330

**Authors:** Julius Durr, Guilhem Reyt, Stijn Spaepen, Sally Hilton, Cathal Meehan, Wu Qi, Takehiro Kamiya, Paulina Flis, Hugh G. Dickinson, Attila Feher, Gary D. Bending, Paul Schulze-Lefert, David Salt, Jose Gutierrez-Marcos

## Abstract

The Casparian Strip (CS) constitutes a physical diffusion barrier to water and nutrients in plant roots, and is formed by the polar deposition of lignin polymer in the endodermis. This precise pattern of lignin deposition is thought to be mediated by the scaffolding activity of membrane-bound Casparian Strip domain proteins (CASPs). However, we show that endodermis-specific receptor-like kinase 1 (ERK1) and ROP Binding Kinase1 (RBK1) are also involved in this intricate process, with the former playing an essential role both in the localization of CASP1 and in lignin deposition. We further characterised ERK1 and determined its subcellular localisation in the cytoplasm and nucleus of the endodermis, as well as provide evidence for its involvement in a signalling pathway together with the circadian clock regulator, Time for Coffee (TIC). We also show that disruption to CS organisation and increased suberisation in the endodermis due to loss of function of either *ERK1* or *TIC* collectively leads to an altered root microbiome composition. Thus, our work reveals additional players in the complex cascade of signalling events operating in the root endodermis to establish both the CS diffusion barrier and the microbial composition of the rhizosphere.

## Introduction

Plant roots are specialised structures essential for plant growth and survival. Roots control the selective uptake of nutrients and water from the soil, and also prevent the passive diffusion and entry of pathogens and toxins (Hawes et al., 1998; Parniske, 2008), while attracting beneficial microorganisms through the secretion of certain compounds into the soil (Sasse et al., 2018). This selectivity is determined by the root architecture and its ability to confer a barrier between the vascular cylinder and the outer cell layers, primarily cortex and epidermis, which are connected to the soil via the apoplast to control the uptake and the exudation of compounds. The apoplastic barrier provided by the Casparian strip (CS) is contained within the endodermis; the innermost cortical cell layer surrounding the vasculature (Bonnett, 1968; Nagahashi et al., 1974). This specialised group of cells possess highly localised ring-like lignin deposits in the radial and transverse cell walls surrounding the endodermis (Naseer et al., 2012). This lignification breaks the apoplastic connection between the vasculature and the outer cell layer by sealing the cell wall.

The endodermal cell wall undergoes further modification by the incorporation of suberin – a hydrophobic polymer composed of long chain fatty acids and glycerol embedded with waxes (Baxter et al., 2009), which constitutes a second barrier. The suberin lamellae ensure that, in fully differentiated endodermal cells, nutrients can only enter or leave the vasculature through specialised transmembrane carriers or special unsuberised passage cells (Graca and Santos, 2007; Doblas et al., 2017a; Andersen et al., 2018). The extent of suberin deposition is dynamic and mediated by abscisic acid and ethylene in response to nutrient stresses (Barberon et al., 2016) or in the case of a defective CS (Wang et al., 2019). Further, suberisation shows high plasticity in contrast to lignification of the CS.

In Arabidopsis, the formation of the CS apoplastic barrier is regulated by the transcription factor MYB36, which controls the expression of major genes involved in CS formation during the initial stages of endodermis differentiation (Kamiya et al., 2015). The first major MYB36-dependent initiation step that gives rise to CS formation is the polar localisation of five redundant CASPARIAN STRIP DOMAIN PROTEINs (CASP1-5) at the site of CS initiation (Roppolo et al., 2011). The CASPs form a platform to recruit proteins involved in polar lignin deposition, such as endodermis specific peroxidase PEROXIDASE64 and NADPH oxidase F (RbohF) (Lee et al., 2013). Several mutants have been identified that affect CASP localisation to the endodermis. The two receptor-like kinase mutants *schengen 1* (*sgn1)* and *schengen 3* (*sgn3*), and the dirigent domain-containing protein mutant *enhanced suberin 1* (*esb1*) exhibit only a discontinuous CS, while *CASP* expression and lignification of the CS domain remain unaffected (Hosmani et al., 2013; Pfister et al., 2014; Alassimone et al., 2016). By contrast, other CS mutants, such as *lord of the rings 1* and *2* (*lotr1* and *lotr2*), display relatively strong ectopic localisation of CASP proteins outside of the CS domain and irregular lignification (Kalmbach et al., 2017; Li et al., 2017).

Despite these findings, the precise mechanisms underlying formation of the endodermal barriers, and in particular CS lignification, remains poorly understood. To further understand these events, we performed a reverse genetic screen to identify novel signalling components involved in endodermis development. Here, we report two cytoplasmic receptor-like kinases that specifically accumulate in the root endodermis and are involved in the formation of the CS apoplastic barrier. We show that these kinases facilitate the polar localisation of CASP1 and the polar deposition of lignin at the CS domain. In addition, we provide evidence that the circadian clock regulator protein, TIME FOR COFFEE (TIC), is a downstream target of these kinases and is also involved in CS organisation. Finally, we also show that the correct deposition of lignin and suberin in the endodermis is critical for the selective recruitment of microbes to roots.

## Results

### Identification of two cytoplasmic receptor-like kinases implicated in the formation of the endodermal barriers

To identify novel components of the signalling pathway implicated in the formation of the CS, we studied the expression of receptor-like kinases (RLK) that are under the control of MYB36 – a transcription factor implicated in endodermis gene expression and CS formation (Kamiya et al., 2015). We identified a candidate, RLK (At5g65530/ ARLCK VI_A3), hereafter named ENDODERMIS RECEPTOR KINASE1 (ERK1) based on the localization of a ERK1-GFP transcriptional fusion in the cytoplasm and nucleus of endodermal root cells only (Fig. 1a). We found that MYB36 was able to bind specifically to a discrete portion of the ERK1 promoter (Fig. 1b) and that the expression of ERK1-GFP is first detected in the late elongation zone and reaches its highest levels in the region where the first xylem bundles are formed (Fig. 1a, Supplementary Fig. 1a). The expression of ROP Binding Kinase1 (RBK1) (Waese et al., 2017), the closest homologue of ERK1, was also restricted to the root endodermis, as confirmed by our transcriptional RBK1-GFP reporter (Supplementary Fig. 1b). Both ERK1 and RBK1 lack the extracellular and transmembrane domains usually present in other receptor-like kinases (Afzal et al., 2008) but have a conserved serine/threonine kinase domain that is activated by phosphorylation through small monomeric G proteins of the plant-specific Rho family (RAC/ROP)(Huesmann et al., 2012). While genetic analyses of ERK1 and RBK1 have previously revealed their roles in trichome branching and in the control of basal resistance to powdery mildew (Huesmann et al., 2012; Reiner et al., 2015), and in auxin-responsive cell expansion (Enders et al., 2017) and pathogen response (Molendijk et al. 2008), respectively, their roles in endodermal function are unknown. We therefore isolated independent T-DNA insertion lines for both genes and assessed mutants for defects in apoplastic barrier diffusion determined by leakage analysis of the apoplastic tracer propidium iodide (PI) into the stele of roots (Supplementary Fig. 1c) (Reiner et al., 2015). We found that all ERK1 mutant lines (*erk1-1*/SALK_148741, *erk1-2*/SALK_010841 and *erk1-3*/SALK_060966) showed defects in the formation of the apoplastic barrier in the endodermis, whereas RBK1 mutants (*rbk1-1*/SALK_043441) did not display any obvious defects (Fig. 1b,c). Intriguingly, *erk1-3*/*rbk1-1* roots showed a significantly stronger defective apoplastic barrier phenotype than either of the individual single mutants, as measured by PI penetration (Fig. 1b,c), suggesting that *rbk1-1* has an additive effect in the absence of *ERK1*. In addition, we were able to restore the apoplastic barrier defects observed in *erk1-3* by complementation with a wild-type *ERK1* genomic fragment (Supplementary Fig. 1d), suggesting that the CS defects observed are due to the loss of *ERK1* function.

**Figure 1.**
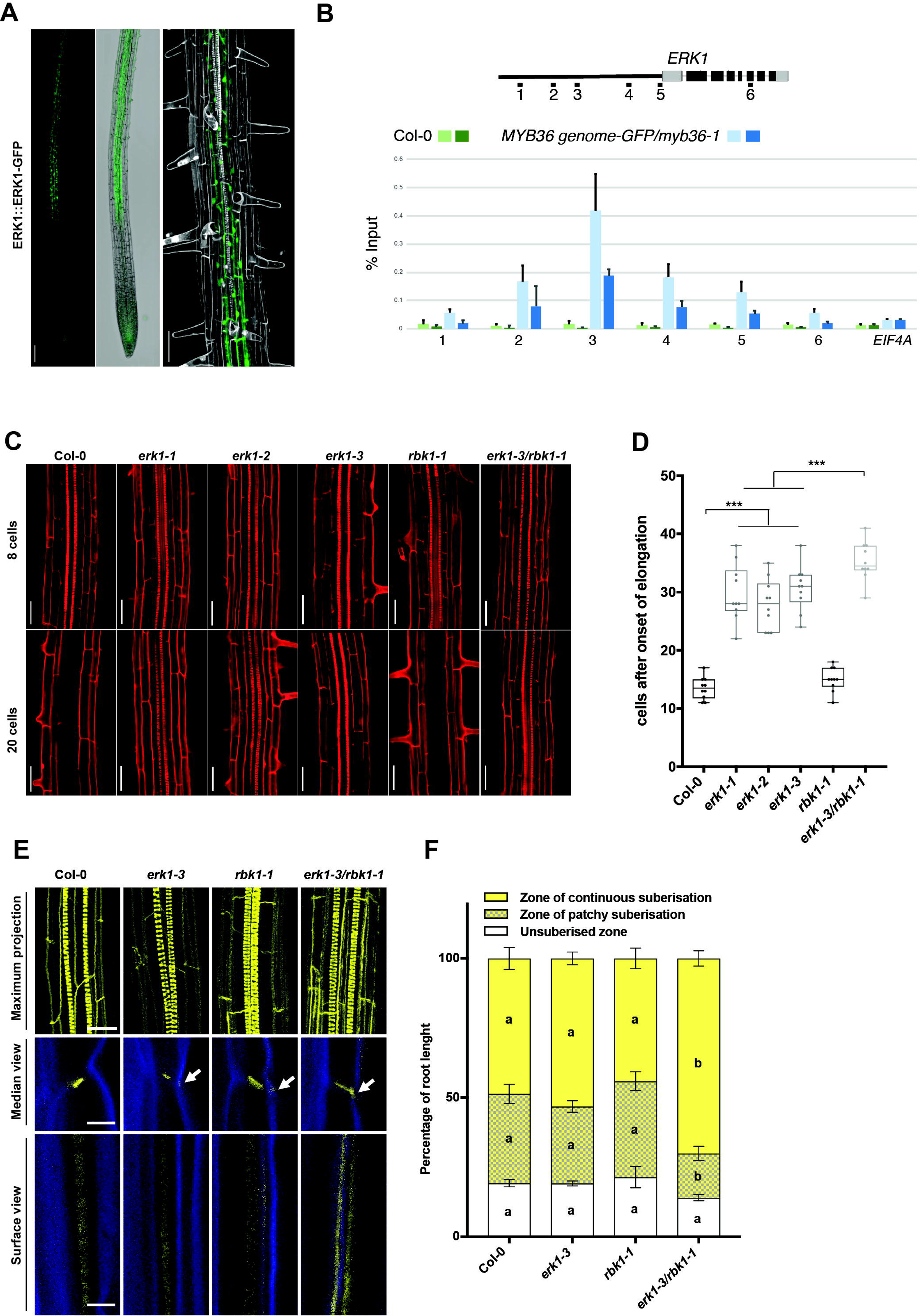
Loss of Casparian strip integrity and disruption of the apoplastic barrier in *erk1* and *rbk1* mutants. A CLSM images of roots expressing ERK1–GFP. Cell walls stained with propidium iodide (grey). Bar = 200 μM B Chromatin immunoprecipitation shows that MYB36 binds the promoter of ERK1. C Lack of endodermal diffusion barrier in *erk1* and *rbk1* mutants visualized by presence of propidium iodide (red) in stele. Bar = 50 μM D Quantification of PI penetration into the stele quantified as number of endodermal cells from the first fully expanded cell (n = 10). E Three-dimensional maximum projections of Casparian strip autofluorescence. Spiral structures in the centre of the root are xylem (top). Longitudinal section of lignin deposition sites (bottom). Cleared roots were stained with basic fuchsin (yellow; lignin) and Calcofluor White (blue; cellulose). Although both of these dyes stain cell walls, basic fuchsin primarily interacts with lignin and Calcofluor White with cellulose. F Quantitative analysis of suberin accumulation. Suberin was stained with fluorol yellow 088. The endodermal cell suberin was counted from the onset of elongation to the junction (base) between root and hypocotyl (n = 6)

We next assessed endodermis cell wall deposition in *erk1* mutants by electron microscopy and found it to be thickened in mutants compared to wild-type roots (Supplementary Fig. 2). To further investigate the defective apoplastic barrier observed in the endodermis of *erk1* mutants, we then examined the pattern of lignin and cellulose deposition in the CS using dyes and confocal microscopy. Compared to wild-type roots, the single *erk1-3* and *rbk1-1* mutants showed a slight accumulation of ectopic lignin in the corners of endodermal cells, though ectopic lignin was more apparent in *erk1-3*/*rbk1-1* mutants (Fig. 1d). The precocious and ectopic deposition of suberin has also been observed in the endodermis of other mutants harbouring similar disruptions to the CS, such as *esb1-1*, *casp1-1/casp3-1*, *myb36-2* and *lcs2-1* (Roppolo et al., 2011; Hosmani et al., 2013; Kamiya et al., 2015; Li et al., 2017) and so assessed suberin content in our mutants. We found that suberisation in single *erk1-3* and *rbk1-1* mutants was normal, while *erk1-3*/*rbk1-1* roots showed premature suberisation (Fig. 1e). Collectively, these data suggest that both ERK1, and to a lesser extent RBK1, affects lignin and suberin deposition in the endodermis.

### Downstream targets of ERK1 are required for CS barrier formation

It has been reported that ERK1 is capable of phosphorylating proteins *in vitro* (Reiner et al., 2015). Therefore, to identify downstream targets of this kinase, we performed a mass spectrometry-based quantitative phosphoproteomics analysis using roots from wild-type (Col-0) and *erk1-3* plants. We identified 100 proteins displaying over 1.5-fold significant change in abundance (Supplementary Fig. 3a and Supplementary Table 1). To reveal the potential targets and pathways affected, we performed a Gene Ontology (GO) analysis and found sequences associated with the terms “response to abscisic acid”, “establishment of localisation” and “translation” to be significantly enriched (p<0.001) (Supplementary Fig. 3b). From this analysis we focused our investigation on two notable proteins, TIME FOR COFFEE (TIC) and TOM-LIKE6 (TOL), which were differentially phosphorylated in *rck1-3* (Supplementary Fig. 3c,d), and have been implicated in different aspects of plant development. To evaluate the possible involvement of these proteins in the formation of the CS barrier, we assessed the permeability of the apoplastic tracer PI into the stele in the respective mutant backgrounds. We observed a large delay in the formation of the PI block in *tic-2* mutants, which was significantly more than in the *erk1-3* mutant allele (Fig 2a). By contrast, we did not see any increased permeability in *tol6-1* or in combinations of other *TOL* mutants (Fig. 2b and Supplementary Fig. 4a). To evaluate if the *tic-2* apoplastic barrier defect is linked to circadian clock defects, we analysed three circadian clock mutants (*elf3-4*, *cca1-1* and *elf4-7)*, however none of these mutants showed any changes with respect to the wild-type (Supplementary Fig. 4b). We then inspected the deposition of lignin in the endodermis of *TIC* and *TOL* mutants and found that both *tic-2* and the quintuple *tolQ* showed strong ectopic deposition of lignin in the lateral margins of endodermal cells (Fig. 2c). Intriguingly, we found that only *tic-2* showed precocious accumulation of suberin in the endodermis (Fig. 2c). Collectively, these results suggest that ERK1 and TIC act in in the same pathway and are responsible for the organisation of the CS. Additionally we looked at PI penetration, lignin deposition and suberin accumulation in mutants of *erk1-3* in combination with *tic-2* and general CS mutants *myb36-2, sgn3-3, esb1-1* and *casp1-3/casp3-1*. We assessed PI penetration and lignin deposition but found no significant additive effects in double *erk1-3/tic2* mutants compared with single *erk1-3* mutants, nor in any of the other double-mutants compared with the single mutants, except for *erk1-3/sgn3-3*, which seemed to contain slightly less lignin than *sgn3-3* (Supplementary Fig. 5 and 6). When assessing suberin accumulation, again no major differences were observed, however, triple and double mutants of *erk1-3* with *casp1-3/casp3-1* and *myb36-2,* respectively, appeared to have slightly less suberin (Supplementary Fig. 7). Two recently identified stele-expressed peptides, CASPARIAN STRIP INTEGRITY FACTORS 1 & 2 (CIF1 & 2), which bind to the SGN3 receptor and promote Casparian strip formation, have been shown to enhance suberin deposition in wild-type plants and induce CS mislocalisation as well as overlignification (Doblas et al., 2017b; Nakayama et al., 2017). To determine whether ERF1 and RBK1 kinases are linked with the CIF/SGN3 signalling pathway we assessed lignin deposition, CASP1-GFP expression and suberin accumulation in response to exogenous application of the CIF peptide. Our data shows that CIF2 induced suberisation, ectopic polymerisation and mislocalisation of CASP1-GFP similar to wild-type, suggesting that the role of ERF1/RBK1 in suberin accumulation is part of an additional compensatory pathway independent to the SGN3/CIF pathway (Supplementary Fig. 8 and 9).

**Figure 2.**
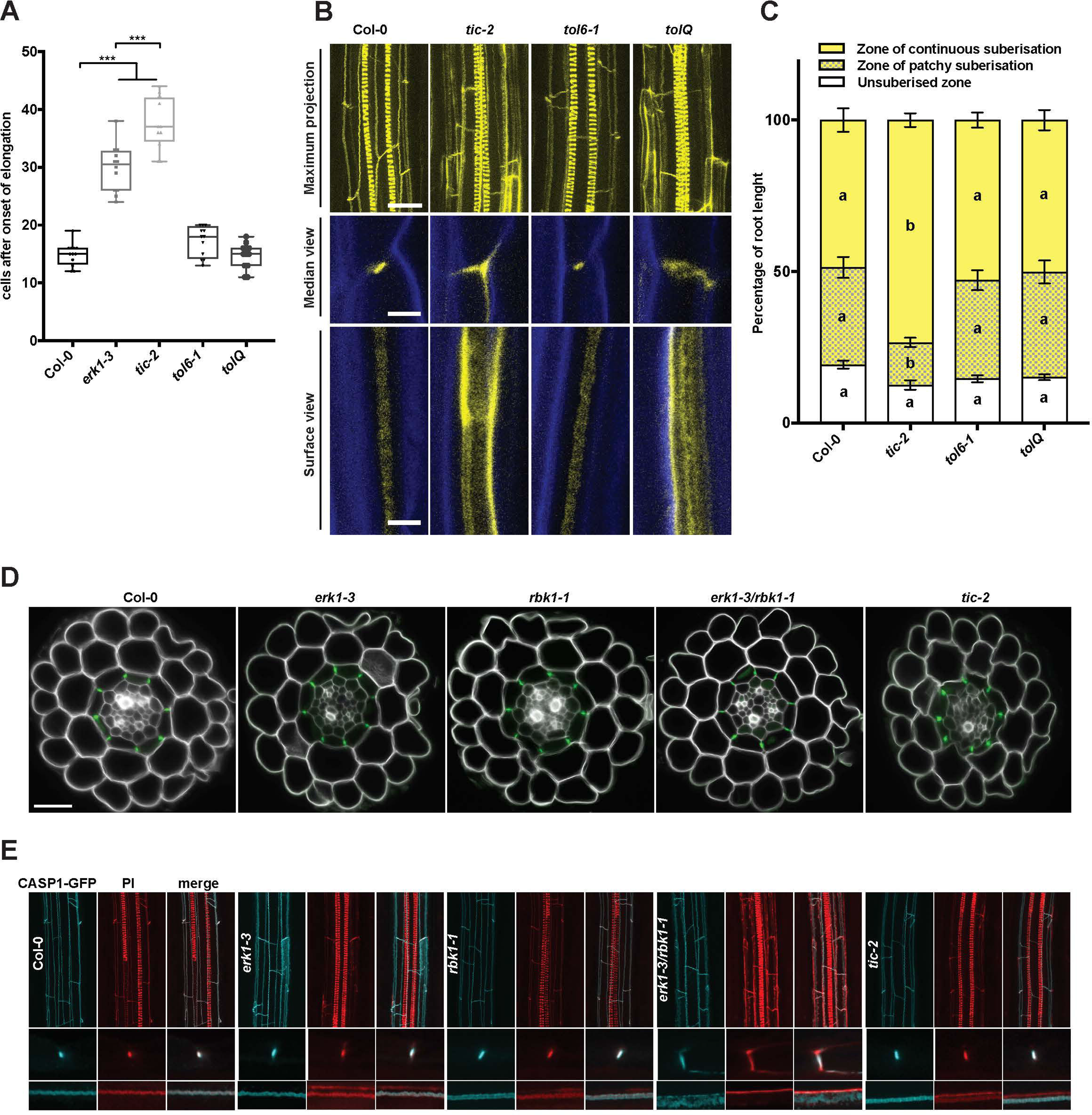
*TIC* and *TOL* are downstream targets of the ERK1 signalling pathway implicated in casparian strip formation. A Quantification of PI penetration into the stele quantified as number of endodermal cells from the first fully expanded cell (n = 10). B Three-dimensional maximum projections of Casparian strip autofluorescence. Spiral structures in the centre of the root are xylem (top). Longitudinal section of lignin deposition sites (bottom). Cleared roots were stained with basic fuchsin (yellow; lignin) and Calcofluor White (blue; cellulose). Although both of these dyes stain cell walls, basic fuchsin primarily interacts with lignin and Calcofluor White with cellulose. C Quantitative analysis of suberin accumulation. Suberin was stained with fluorol yellow 088. The endodermal cell with suberin was counted from the onset of elongation to the junction (base) between root and hypocotyl (n = 6). D CLSM images of cross-section from roots expressing CASP1–GFP. Cell walls stained with propidium iodide (grey). Bar = 20 μM E Three-dimensional maximum projections of the mature endodermis expressing CASP1–GFP and stained with basic fuchsin (lignin, red) (top) on cleared roots. Median and surface view of mature endodermal cells expressing CASP1–GFP and stained with basic fuchsin (bottom).

The ectopic deposition of lignin in *erk1-3/rbk1-1* and *tic-2* mutants raises the possibility that the CS machinery is mislocalised in these mutants. To evaluate this hypothesis, we assessed the localisation of the CASP1-GFP reporter in the different mutant backgrounds. We did not observe any major defects in the cellular organisation of root optical cross-sections in any of the mutants tested (Figure 2b). However, in transverse root cross-sections, we noticed that the polar localisation of CASP1-GFP and the deposition of lignin was altered in the CS domain of *erk1-3* /*rbk1-1* mutants (Fig. 2d,e). Taken together, these data suggest that ERK1 and RBK1 affect the polar localisation of CASP1 in the endodermis.

### The endodermal defects in *erk1erk1-3 and tic-2* lead to ionomic changes

Root suberisation and the CS have been shown to play a role in environmental adaptation by acting as a physical barriers to prevent unfavourable inward and outward leakage of ions between the xylem and the soil (Barberon et al., 2016). In previous studies, it was shown that several CS-defective mutants exhibit changes in concentrations of multiple elements. Therefore, we compared the ionomic profiles of *erk1-3, rbk1-*1 and *tic-2 mutants* with other mutants known to be defective in CS function. A principal-component analysis of elemental concentrations in the leaves showed that the ionomic profile of the *erk1-3* and *rbk1-*1 mutants was similar to wild-type, whereas the *erk1-3/rbk1-*1 double mutant was more similar to *esb1-1* but distinct to either *myb36-2* or *sgn3-3* mutants (Supplementary Fig. 10a). Unlike *sgn3-3, esb1-1 and myb36-2* have enhanced lignification and suberisation (Hosmani et al., 2013). A more detailed analysis of the ionomic data revealed that *erk1-3/rbk1-*1 mutants accumulated higher levels of iron in their roots compared with wild-type plants (Fig 3a). Because iron is more soluble in acidic soils than in neutral soils, we reasoned that this response may reflect an adaptation to ensure the growth and survival of plants under unfavourable mineral conditions. We therefore tested the effect of iron on ERK1-GFP accumulation and found that ERK1-GFP expression in the endodermis increased in response to excessive iron in a pH-dependant manner (Fig 3d). Because a previous genetic screen for iron homeostasis mutants identified TIC as a regulator of FERRITIN1 (Duc et al.,2009) we tested the growth response of these mutants under high iron growth conditions. We found that both *erk1-3* and *tic-2* displayed high sensitivity to elevated iron conditions (Fig. 3b,c) as well as defects in germination under other ion stress conditions (Supplementary Fig. 10b-d). High sensitivity to iron was also observed in *sgn3-3* but not in *myb36-2* (Supplementary Fig. 11). Collectively, these results suggest that these mutants are affected in their responses to fluctuating iron levels most likely due to the inward and outward leakage resulting from their defective endodermal barriers.

**Figure 3.**
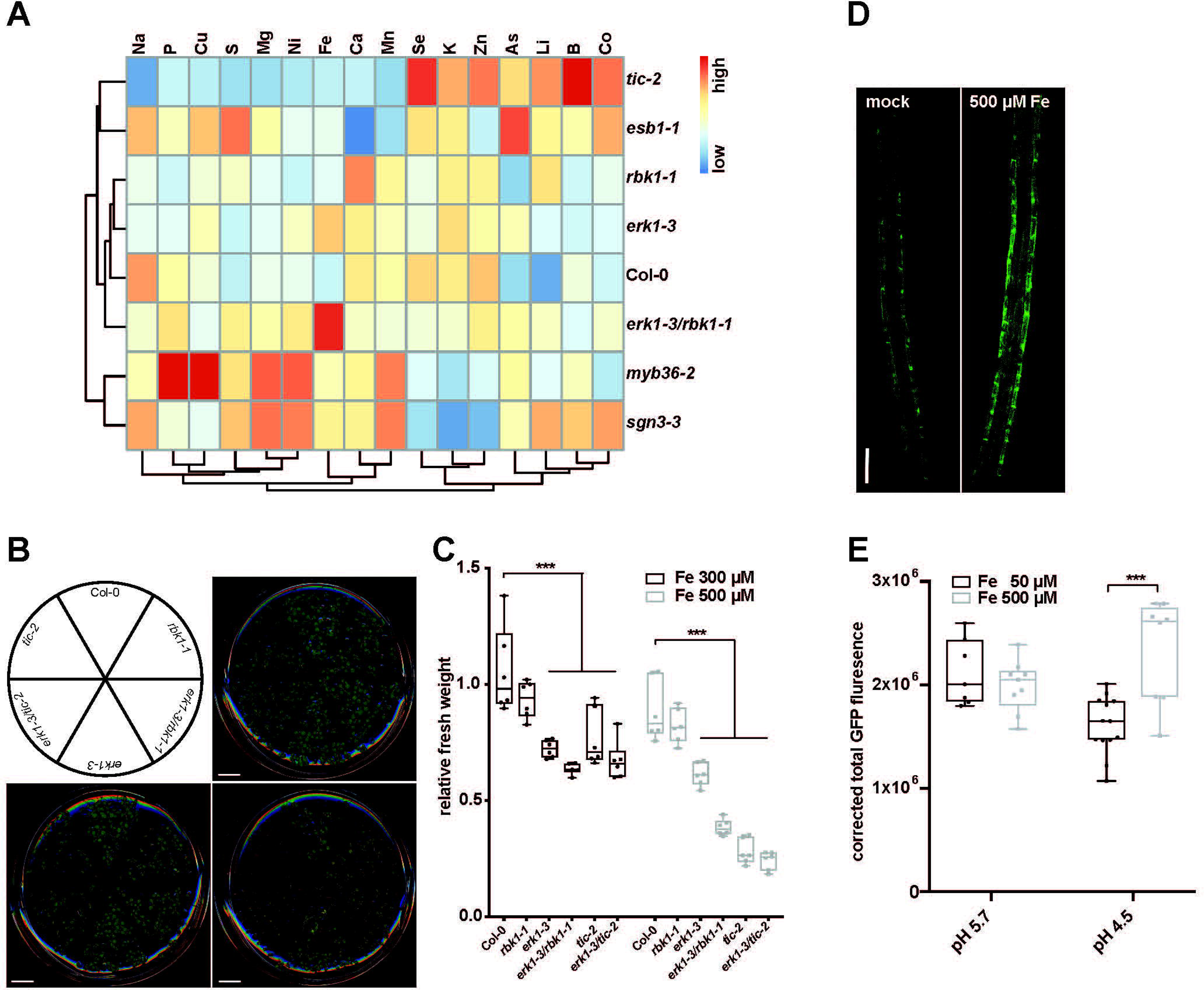
ERK1 and TIC mutants display ionomic changes and are sensitivity to excess iron. A Heat-map ionomic analysis of shoots from wild-type and mutants (n = 15). B Iron sensitivity of plants grown *in vitro* under different iron concentrations. Media containing 50 μM iron-EDTA (control; top right), Media containing 300 μM iron-EDTA (bottom left) or Media containing 500 μM iron-EDTA (bottom right). C Relative mean fresh weight compared to the control (n=6). The plants were grown as described for B. D Induction of ERK1-GFP expression in root endodermis by elevated iron-EDTA. E Quantification of ERK1-GFP expression in endodermis by iron-EDTA at different pH.

### The root endodermal barrier influences the structure of the root microbial community

It has previously been shown that altered exudation from the plant root causes changes in the composition of the rhizosphere microbiome (Sasse et al., 2018). On this basis, we reasoned that the unregulated leakage of components into and out of *erk1-3* and *tic-2* mutants might also affect their respective root microbiomes. To test this hypothesis, we grew wild-type and single mutant plants in natural soils, and after four weeks we analysed the bacterial communities present in the root rhizosphere. We found clear differences between the genotypes present in each subpopulation, where the communities associated with *erk1-3* and *tic-2* mutants were most similar but differed significantly from wild-type plants (Fig. 4a). The results were further validated by an analysis of similarities (ANOSIM), which revealed significant variations in the microbial communities from roots of wild-type plants and *erk1-3* (ANOSIM, r=0.921, p=0.001) or *tic-2* plants (ANOSIM, r=0.922, p=0.001) (Fig. 4b and Supplementary Fig. 12a), whereas no significant differences were observed between the *erk1-3* and *tic-2* microbiomes (ANOSIM, r=-0.019, p=0.558).

**Figure 4.**
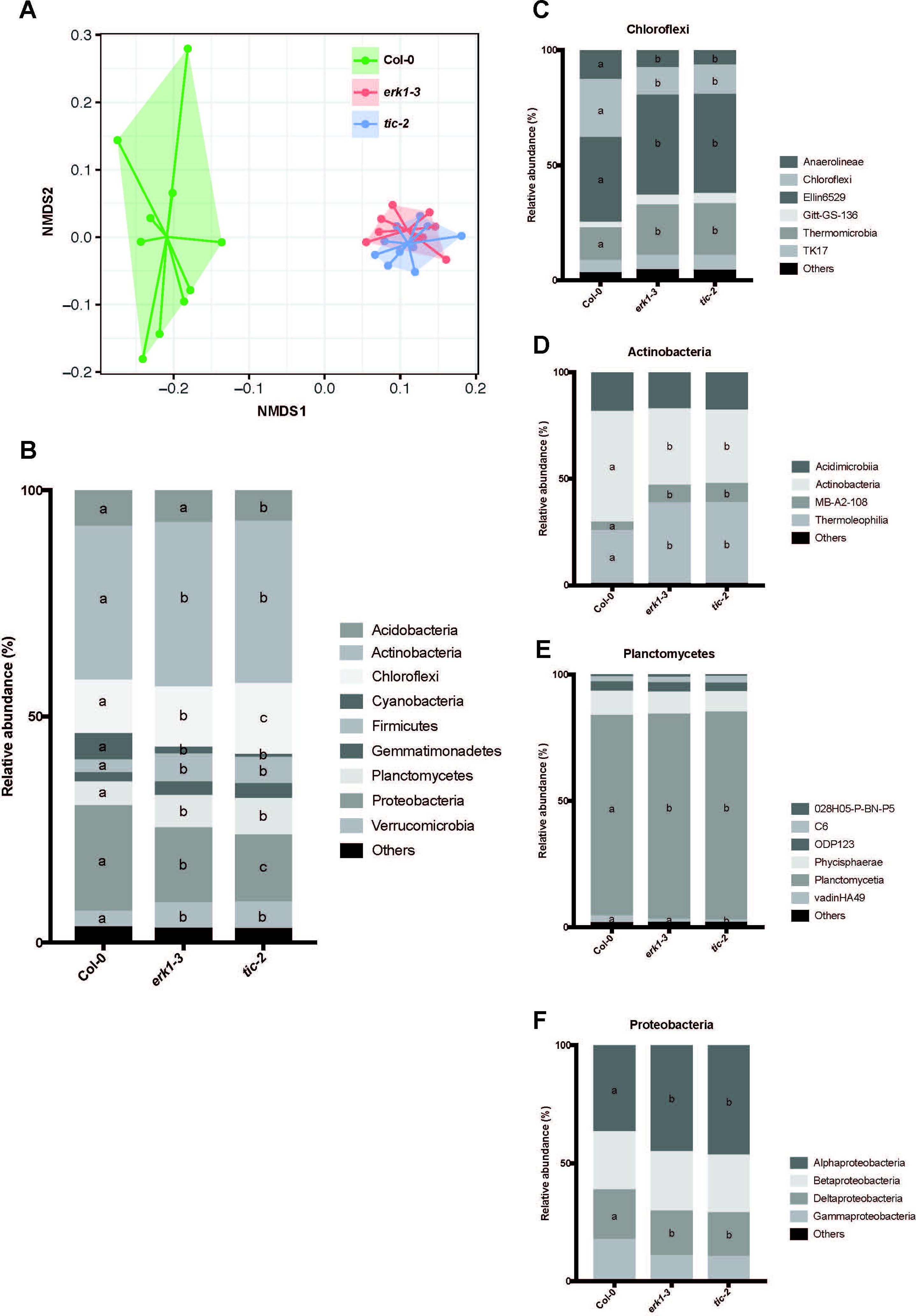
Differences in microbial community present in *erk1-3* and *tic-2*. A PCA of Bray-Curtis distances of bacterial communities present in roots *erk1-3* and *tic-2* plants grown in natural soils (n=10). B Distribution of soil bacteria in roots of Col-0 (WT), *erk1-3* and *tic-2* plants. plants over two consecutive generations (G2 and G3). C-F Distribution of Firmicutes, Chloroflexi, Actinobacteria, Planctomycetes and Proteobacteria in roots of Col-0 (WT), *erk1-3* and *tic-2* plants.

We next carefully examined the different phyla of root-associated microbes and identified several groups that differed in abundance between the *erk1-3* and *tic-2*-associated populations (Fig. 4c). Notably, Cloroflexi and Proteobacteria populations differed significantly amongst the three genotypes tested (Fig. 4d-f). To independently validate these results, we inoculated seeds with a synthetic bacterial community (SynCom) isolated from an Arabidopsis root rhizosphere (Bai et al., 2015) and determined the microbial communities present in mature roots and leaves. We found differences between the microbiomes present in leaves (25.4% of variance in leaves explained by genotype; permutation-based ANOVA test p=0.002) and roots (10.2% of the variance explained by genotype; permutation-based ANOVA test, p=0.27) of wild-type and the two mutants (Supplementary Fig. 12a,b). Most of the differences in leaves were associated with the abundance of Xanthomonadaceae (*Proteobacteria*) and *Flavobacteriaceae* (*Bacteroidetes*) (Supplementary Fig. 12c). Taken together, our results indicate that the capacity of microbes to colonize roots is strongly influenced by lignification and suberisation of the endodermal barriers.

## Discussion

The polarised deposition of cell wall material is crucial for the function and development of many cell types (Roppolo and Geldner, 2012). One of the most-studied examples of polar cell wall deposition is the endodermis and its ring-like CS that prevents the inward and outward leakage of metabolites from the vasculature (Doblas et al., 2017a). One of the critical steps in the separation of the inside and outside facing plasma membrane domains is the proper deposition of CASP proteins to the site of lignin deposition (Roppolo et al., 2011). Various mutants with defective CSs have been described in recent years. In addition to the CS defects, most of these mutants also show a mis-localisation of CASP proteins at the site of lignification. Three different classes of CS mutants have been identified according to CASP1 expression and localisation: i) *MYB36,* is a key determinant of CS formation and its loss-of-function mutant *myb36-3*, shows no CASP1-GFP expression; ii) mutations such as *sgn1*, *sgn3* and *esb1* do not affect the central positioning of CASPs, only the continuity of the CASP domain (Hosmani et al., 2013; Pfister et al., 2014; Alassimone et al., 2016); iii) mutants such as *lotr1/lcs2* or *lotr2*/exo70a1 exhibit ectopic deposition of CASP proteins outside of the normal site of CS lignin deposition (Kalmbach et al., 2017; Li et al., 2017).

In this study we describe mutations affecting the cytoplasmic receptor kinases ERK1 and RBK1, which lead to the mislocalisation of CASP1-GFP and ectopic lignin deposition outside of the CS. Further, the lack of a proper apoplastic barrier in the double mutants is accompanied by an increase in lignin and suberin, which has been attributed to a compensatory mechanism to counteract the loss of an apoplastic barrier in the endodermis (Doblas et al., 2017a). This effect is likely dependant on the membrane-bound and cytoplasmic kinase module SNG3/SGN1 (Pfister et al., 2014; Alassimone et al., 2016). The mechanism by which the ERK/RBK1 signalling module affects lignin deposition and suberisation is currently unclear, however, evidence from the literature suggests that these proteins could be involved in the rearrangement of microtubules and membrane trafficking (Huesmann et al., 2012). Indeed, it has been reported that ERK1 mutants exhibit increased trichome branch numbers (Reiner et al., 2015), which is strongly dependent on the correct rearrangement of microtubules (Mathur et al., 1999). Similarly, a transient knockdown of the barley *RBK1* homolog, *HvRBK1,* leads to defects on cortical microtubule stability in epidermal cells (Huesmann et al., 2012). Microtubules have also been shown to play an important role in the polar deposition of subcellular components and proteins that are involved in cell wall formation (Marchant, 1979). EXO70A1, which is important for the polar positioning of CASP proteins at the site of CS formation, together with other members of the exocyst complex, has also been shown to be deposited at the site of secondary cell wall thickenings in tracheary elements, in a microtubule-dependent manner (Kalmbach et al., 2017; Vukasinovic et al., 2017).

In addition to identifying the involvement of ERK1 in CS formation, we also identified TOL6 and TIC as downstream components of this ERK1-mediated signalling pathway. TOL6 is part of the plant Endosomal Sorting Complex Required for Transport (ESCRT) complex (Moulinier-Anzola et al., 2014; Sauer and Friml, 2014) and a member of a gene family that acts redundantly to control plant morphogenesis (Korbei et al., 2013). *TOL6* mutants did not show obvious CS apoplastic barrier defects, however the quintuple mutant (*tol2-1/tol3-1/tol5-1/tol6-1/tol9-1*; *tolQ*) exhibited strong ectopic lignin deposition similar to *erk1-3/rbk1-1* mutants. The ectopic lignification observed in *tolQ* did not interfere with apoplastic barrier function, suggesting that lignin deposition in this mutant is probably a secondary effect of the disruption to vesicle transport. This idea is supported by the fact that in addition to microtubules, membrane vesicle transport also plays a critical role in the localisation of plant cell wall components (Miao and Liu, 2010; McFarlane et al., 2013). TIC is one of the main regulators of the circadian clock (Davis and Millar, 2001; Sanchez-Villarreal et al., 2013) and has been shown to be an integral component of iron homeostasis in roots (Duc et al., 2009; Sanchez-Villarreal et al., 2013). Our analysis revealed that although *tic1-2* mutants show apoplastic barrier defects and ectopic deposition of lignin in the CS, this did not interfere with the polar localisation of CASP1 in the endodermis. TIC is a major regulator of the circadian clock (Davis and Millar, 2001; Sanchez-Villarreal et al., 2013), yet the CS function defects observed in *tic-2* are not a direct cause of circadian rhythm defects. TIC has also been implicated in carbon homeostasis, particularly in sugar production or its responses (Sanchez-Villarreal et al., 2013), both of which are critical factors in cell wall formation. The striking resitance to drought observed in *tic1*-2 mutants (Sanchez-Villarreal et al., 2013) could be explained by an enhanced suberisation of the endodermis that has also been shown in other casparian strip mutants (Baxter et al., 2009).

Plant roots secrete sugars and other metabolites into the soil as a means of attracting beneficial microbes and in defence against pathogens, which ultimately shapes the microbial communities present in the rhizosphere (Lanoue et al., 2010; Sasse et al., 2018). Root exudates are highly variable and defined by many parameters including plant accession (Monchgesang et al., 2016b; Monchgesang et al., 2016a), developmental stage, abiotic stresses (Carvalhais et al., 2013; Chaparro et al., 2014), and more recently by the circadian clock (Hubbard et al., 2018). Given that the CS and suberin lamellae are regulators of water and nutrient uptake in the root endodermis, it is plausible that perturbations in these structures could also shape the composition of the root microbiome by altering exudate secretion, as observed in this study. Indeed, we showed that *erk1-3* and *tic-2* recruit similar microbial populations to their rhizosphere that differ markedly to the wild-type. Alternatively, modifications in the rhizosphere microbiome could be caused by differences in actively secreted compounds associated with suberisation. Suberin lamellae are not thought to affect the apoplastic barrier directly but instead modulate transport through the endodermis (Robbins et al., 2014; Andersen et al., 2015), while some suberin components display antimicrobial properties (Lulai and Corsini, 1998; Thomas et al., 2007; Ranathunge et al., 2008; Buskila et al., 2011). Thus, the increased suberisation observed in *erk1-3* and *tic-2* could account for the specific microbial communities present in these mutant roots.

Taken together our work provides evidence for a complex signalling cascade taking place at the root endodermis, which is necessary for both the formation of a functional CS diffusion barrier and for correct accumulation of suberin, which ultimately determines the microbial composition of the rhizosphere.

## Material and Methods

### Plant lines and growth conditions

All plant material used in this study was derived from the wild-type Columbia (Col-0) or Wassilewskija (Ws) accession. T-DNA insertion alleles of ERK1 (*erk1-1*, *rlck vi_a3-1*: SALK_148741; *erk1-2*, *rlck vi_a3-2*: SALK_010841 and *erk1-3*, *rlck vi_a3-3*: SALK_060966) and RBK1 (*rbk1-1*: SALK_043441) were obtained from the Nottingham *Arabidopsis* Stock Centre (NASC). Mutant and wild-type (Col-0 or Ws) seeds were sown on soil (John Innes and Perlite mix), stratified for 2 days at 4°C in the dark, and grown at 20°C under long photoperiod conditions (150 μmol/m2/s and 16 h/8 h light/dark cycles) to induce flowering.

Seeds were surface sterilized for 3 min in 70% ethanol, followed by a 2 min treatment in 1% NaOCl. Seeds were then washed six times in sterile H_2_O, dispersed in sterile 0.1% agarose and placed on half-strength MS medium (Murashige & Skoog, Sigma), solidified with 0.8% Phytagar. Seedlings were grown vertically in growth chambers at 22°C, under long days (16-hr light/8-hr dark), 100 μE light, and were used at 5 days after shift to room temperature.

For the evaluation of growth effect of excess iron, plants were grown on medium containing 5 mM KNO_3_, 2.5 mM KPO_4_, 3 mM MgSO_4_, 3 mM Ca(NO_3_)_2_, 70 μM H_3_BO_3_, 14 μM MnCl_2_, 0.5 μM CuSO_4_, 1 μM ZnSO_4_, 0.2 μM Na_2_MoO_4_, 10 μM NaCl, 0.01 μM CoCl_2_ and 50 μM, 300 μM or 500 μM Fe-EDTA, solidified with 0.8% phytoagar. Around 30 plants per line were grown on each plate. After 14 days the fresh weight of all plants of one line per plate was measured. Relative fresh weight was calculated by normalisation of the iron excess samples (300 μM and 500 μM Fe-EDTA) by the mock control (50 μM Fe-EDTA).

### Plant transformation

Transgenic plants were generated by introduction of the plant expression constructs into an Agrobacterium tumefaciens strain GV3101 and transformation was done by floral dipping (Clough and Bent, 1998).

### Vector construction

For generation of expression constructs, Gateway Cloning Technology (Invitrogen) was used. pERK1::ERK1 was cloned using the genomic sequence of ERK1 including the endogenous promoter consisting of 1403 bp upstream of the ATG. pERK1::ERK1 was cloned into pGWB440 and into pFAST-R01 and transformed in transformed in Col-0 and *erk1-3* background, respectively. pRBK1::RBK1 was cloned using the genomic sequence of RBK1 including the endogenous promoter consisting of 1996 bp upstream of the ATG. pRBK1::RBK1 was cloned into pGWB440, pGWB553 and into pFAST-R07 and transformed in in Col-0 background.

### Chromatin Immunoprecipitation

Chromatin Immunoprecipitation (ChIP) analysis was performed by following the protocol as described (Nakamichi et al., 2010) with modifications. Roots (100 mg fresh weight) from 11-d-old plants were cross-linked by using 4 mL of the buffer (10 mM PBS, pH 7.0, 50 mM NaCl, 0.1 M sucrose, and 1% formaldehyde) for 1 h at room temperature with the application of three cycles of vacuum infiltration (10 min under vacuum and 10 min of vacuum release). Glycine was added to a final concentration of 0.1 M to stop the cross-linking reaction, and the samples were incubated for a further 10 min. After being washed with tap water, the samples were ground to a fine powder by using a Multibeads Shocker (Yasui Kikai) at 1,500 rpm for 30 sec. The powder was suspended with 2 mL of Lysis buffer [50 mM Tris·HCl, pH 7.5, 100 mM NaCl, 1% Triton X-100, 1 mM EDTA, EDTA-free Complete protease inhibitor (Roche)] and sonicated by using a Bioruptor UCD-250 (Cosmo Bio) with the following setting: mild intensity, 45 cycles (30 s ON and 30 s OFF) at 4 °C. A 100-μL sample of the chromatin sheared to between 200 and 1,500 bp was stored as the input fraction, and the rest (1.9 mL) was mixed with Dynabeads Protein G (Life Technologies) bound with anti-GFP antibody (ab290; Abcam) and incubated for 2 h at 4 °C. The beads were washed with Lysis buffer, twice with high-salt buffer [50 mM Tris·HCl, pH 7.5, 400 mM NaCl, 1% Triton X-100, 1 mM EDTA, and EDTA-free Complete protease inhibitor (Roche)], and then with Lysis buffer. After Elution buffer (50 mM Tris·HCl, pH 8.0, 10 mM EDTA, and 1% SDS) and proteinase K (0.5 mg/mL) were added to the beads, the beads were incubated overnight at 65 °C. The DNA was purified with NucleoSpin Gel and PCR Clean-up (MachereyNagel) with Buffer NTB (Macherey-Nagel). Eluted solutions were used for qPCR. EIF4A (At3g13920) was used as a negative control (Supplementary Table 2). Two independent experiments were performed with three biological replicates for each.

### Permeability of the apoplastic barrier

For the visualisation of the penetration of the apoplastic barrier by propidium iodide, seedlings were incubated in the dark for 10 min in a fresh solution of 15 mM (10 mg/ml) Propidium Iodide (PI) and rinsed two times in water. The penetration of the apoplastic barrier was quantified by the number of cells from the ‘onset of elongation’ until the PI signal was blocked by the endodermis from entering the vasculature. The ‘onset of elongation’ was defined as the point where an endodermal cell in a median optical section was clearly more than twice its width (Alassimone et al., 2010).

### Histological analysis

Tissue was fixed and stained, as described previously (Musielak et al., 2016). In brief, five-day-old seedlings were fixed in 4% para-formaldehyde (PFA) and cell walls were stained with SCRI Renaissance 2200 (SR2200) (0.1% (v/v) SR2200, 1% (v/v) DMSO, 0.05% (v/v) Triton-X 100, 5% (v/v) glycerol and 4% (w/v) para-formaldehyde in PBS buffer (pH 7.4)) in PBS for 15 min at room temperature with no vacuum applied. Fixed seedlings were washed twice with PBS (pH 7.4) and cleared with ClearSee (10% xylitol (w/v), 15% sodium deoxycholate (w/v) and 25% urea (w/v) in water) for 4 days at room temperature (Kurihara et al., 2015). Cleared samples were washed twice with PBS (pH 7.4) and embedded in 4% agarose. The agarose blocks were cut in 200 μm sections with a VIBRATOME® Series 1000 Sectioning System.

### Electron microscopy analysis

For light Transmission electron microscopy (TEM), roots were dehydrated in an ethanol/propylene oxide series, embedded in Spurr’s resin (Premix Kit-Hard, TAAB Laboratory and Microscopy, Aldermaston, UK) and polymerized at 70 °C overnight. Longitudinal ultrathin sections (60 nm) were cut using an ultramicrotome (ultra-RMC Products), mounted on copper grids, and contrasted with 1 % uranyl acetate in water for 25 min followed by lead citrate for 3 min. Sections were examined in a transmission electron microscope (Spirit Biotwin 12 FEI Company). Measurements were carried out using the TEM Imaging Platform program.

### GFP fluorescence measurement

The confocal images of the pERK1::ERK1-GFP reporter under different iron concentrations were taken with a Zeiss Axio Observer.Z1 inverted microscope equipped with a confocal laser-scanning module (Zeiss LSM 880). The exact same settings were used during one experiment. To be able to compare different roots optical median sections were made. The florescent area was selected with FIJI (ImageJ) and the area, area integrated intensity and mean grey value of each image were measured. Intensities were corrected with the mean intensity of areas without signal (background). The corrected total fluorescence (CTF) was calculated by using this formula: CTF=integrated density-(selected fluorescent area*mean background fluorescence).

### Confocal microscopy

Confocal laser scanning microscopy experiments were performed using a Zeiss Axio Observer.Z1 inverted microscope equipped with a confocal laser-scanning module (Zeiss LSM 710 and Zeiss LSM 880, Warwick) or a Leica SP5 and SP8 (Nottingham).

Excitation and emission setting were used as followed: GFP 488 nm, 500 – 550 nm; propidium iodide 516 nm, 560 – 700 nm; tagRFP 561 nm, 578 – 700 nm; SR2200 405 nm, 410 – 550 nm; Calcofluor white 405 nm, 425 – 475 nm; basic fuchsin 561 nm, 570 – 650 nm. For examining CASP1 expression, basic fuchsin and calclofluor white M2R staining, 6-day-old roots were fixed in paraformaldehyde and cleared in ClearSee as described (Ursache et al., 2018). Fluorol yellow staining for visualization of suberin was performed and quantified as described in Naseer et al., 2012 and Barberon et al., 2016, using a fluorescent microscope Leica DM 5000. Confocal images were analysed and contrast and brightness were adjusted with the FIJI package of ImageJ (http://fiji.sc/Fiji) and Adobe Photoshop Elements Editor.

### Plant total protein purification

For the purification of total proteins from roots seedlings were grown hydroponically in phytatrays (Sigma) on a nylon filter (250 μm mesh; NITEX) which allows the roots to grow through into the half-strength MS medium supplemented with 1% (w/v) sucrose (pH 5.7 with KOH)(Bargmann and Birnbaum, 2010). After two weeks roots were harvested and flash-frozen in liquid nitrogen. The root tissue was homogenised with a mortar and pestle cooled with liquid nitrogen. Protein were extracted by adding twice the volume of extraction buffer (50 mM HEPES, 150 mM NaCl, 1 mM EDTA, 20 mM NaF, 1 mM Na2MoO4, 1% (w/v) NP-40, 1 mM PMSF, 2 μM Calyculin A, 1 mM NaVO4, 1 mM DTT, Protease inhibitor cocktail (Roche) and 2% (w/v) PVPP) to 3 g of root tissue. After 30 min the samples were spun for 15 min at 4,000 g (4°C) to remove debris. The supernatant was transferred to a new tube and centrifuged for another 30 min at 16,000 g (4°C). The supernatant was again transferred to a new tube and methanol/chloroform precipitation was carried out by adding 4 volumes of methanol, 1 volume of chloroform, and 3 volumes of water, respectively. Samples were centrifuges for 15 min at 4000 g and the aqueous top phase was removed. The proteins were precipitated by adding another 4 volumes of methanol and centrifugation for 15 min at 4000 g. The pellet was washed twice with methanol and resuspended in 25 mM HEPES (pH 8). Reduction and alkylation of the cysteine residues was carried out by adding a combination of 5 mM tris(2-carboxyethyl)phosphine (TCEP) and 10 mM iodoacetamide (IAA) for 30 min at room temperature in the dark. The protein was digested with trypsin (Promega Trypsin Gold, mass spectrometry grade) overnight at 37 °C at an enzyme-to-substrate ratio of 1:100 (w:w). After the digested the peptides were suspended in 80% acetonitrile (AcN), 5% trifluoroacetic acid (TFA) and the insoluble matter was spun down at 4000 g for 10 min. The supernatant was used the enrichment of phosphopeptides.

### Phosphopeptide enrichment

The enrichment of phosphopeptides was carried out as described previously with minor modifications (Thingholm et al., 2006). The peptide concentration was measured with a Qubit™ fluorometer (Invitrogen) and 1 μg total peptides were used for each sample. The Titansphere TiO_2_ 10 μm beads (GL Sciences Inc.) were equilibrated in a buffer containing 20 mg/mL 2,5-dihydroxybenzoic acid (DHB), 80% ACN and 5% TFA in a ratio of 10 μl DHBeq per 1 mg beads for 10 min with gentle shaking at 600 rpm. TiO2 beads were used in a ratio of 1:2 peptide-bead ration (w:w). The TiO2 solution was added to each sample and incubated for 60 min at room temperature. The samples were spun down at 3000 g for 2 min and resuspended in 100 μL Wash buffer I (10% AcN, 5% TFA). The resuspended beads were added to self-made C8-columns. C8-colums were made of 200 μL pipette tips with 2 mm Empore™Octyl C8 (Supelco) discs. The columns were spun down at 2600 g for 2 min, washed with 100 μL Wash buffer II (40% AcN, 5% TFA) and 100 μL Wash buffer III (40% AcN, 5% TFA). The peptides were eluted from the TiO_2_ beads with 20 μL 5% ammonium hydroxide and subsequently with 20 μL 20% ammonium hydroxide in 25% AcN. Both eluates were pooled, the volume was reduced to 5 μL in a centrifugal evaporator (20 – 30 min) and acidified with 100 μL of buffer A (2% AcN, 1% TFA). Samples were desalted with a self-made C18 column (Empore™Octadecyl C18). C18 were made in the same way as the C8-columns. Before adding the samples, the C18-columns were activated with 50 μL methanol and washed with 50 μL AcN and 50 μL buffer A* (2% AcN, 0.1% TFA). Samples were loaded on the C18-column and spun at 2000 g for 7 min. The columns were washed with 50 μL ethyl acetate and 50 μL buffer A* and then eluted consecutively with 20 μL 40% AcN and 20 μL 60% AcN. Samples were then vacuum-dried and prior to MS analysis resuspended in 50 μL buffer A*.

### Mass spectrometry

Reversed phase chromatography was used to separate tryptic peptides prior to mass spectrometric analysis. Two columns were utilised, an Acclaim PepMap μ-precolumn cartridge 300 μm i.d. × 5 mm 5 μm 100 Å and an Acclaim PepMap RSLC 75 μm × 25 cm 2 μm 100 Å (Thermo Scientific). The columns were installed on an Ultimate 3000 RSLCnano system (Dionex). Mobile phase buffer A was composed of 0.1% formic acid in water and mobile phase B 0.1 % formic acid in acetonitrile. Samples were loaded onto the μ-precolumn equilibrated in 2% aqueous acetonitrile containing 0.1% trifluoroacetic acid for 8 min at 10 μL min-1 after which peptides were eluted onto the analytical column at 300 nL min-1 by increasing the mobile phase B concentration from 3% B to 35% over 40 min and then to 90% B over 4 min, followed by a 15 min re-equilibration at 3% B.

Eluting peptides were converted to gas-phase ions by means of electrospray ionization and analysed on a Thermo Orbitrap Fusion (Q-OT-qIT, Thermo Scientific). Survey scans of peptide precursors from 350 to 1500 m/z were performed at 120K resolution (at 200 m/z) with a 4 × 105 ion count target. Tandem MS was performed by isolation at 1.6 Th using the quadrupole, HCD fragmentation with normalized collision energy of 35, and rapid scan MS analysis in the ion trap. The MS2 ion count target was set to 1×104 and the max injection time was 200 ms. Precursors with charge state 2–7 were selected and sampled for MS2. The dynamic exclusion duration was set to 45 s with a 10 ppm tolerance around the selected precursor and its isotopes. Monoisotopic precursor selection was turned on. The instrument was run in top speed mode with 2 s cycles.

### Data Analysis

A label-free peptide relative quantification analysis was performed in Progenesis QI for Proteomics (Nonlinear Dynamics, Durham). To identify peptides, peak lists were created by using Progenesis QI. The raw data was searched against Arabidopsis TAIR database and the common Repository of Adventitious Proteins (http://www.thegpm.org/cRAP/index.html) using uninterpreted MS/MS ions searches within Mascot engine (http://www.matrixscience.com/). Peptides were generated from a tryptic digestion with up to two missed cleavages, carbamidomethylation of cysteines as fixed modifications, oxidation of methionine and phosphorylation of serine, threonine and tyrosine as variable modifications. Precursor mass tolerance was 5 ppm and product ions were searched at 0.8 Da tolerances.

Scaffold (TM, version 4.4.5, Proteome Software Inc.) was used to validate MS/MS based peptide and protein identifications. Peptide identifications were accepted if they could be established at greater than 95.0% probability by the Scaffold Local FDR algorithm. Protein identifications were accepted if they could be established at greater than 99.0% probability and contained at least one identified peptide. Proteins that contained similar peptides and could not be differentiated based on MS/MS analysis alone were grouped to satisfy the principles of parsimony.

### Ionome analysis

Ionomic analysis of plants grown liquid media agar was performed as described in Hosmani et al., 2013. Plants were grown on liquid medium (0.25 mM CaCl2 2 H2O, 1 mM KH2PO4, 0.05mM KCl, 0.25 K2SO4, 1 mM MgSO4 7 H2O, 0.1 mM NaFe-EDTA, 2 mM NH4NO3, 30 μM H3BO3, 5 μM MnSO4 5 H2O, 1 μM ZnSO4 7 H2O, 1 μM CuSO4 5 H2O, 0.7 μM NaMoO4 2 H2O, 1 μM NiSO4 6 H2O) in short day condition (10h light 20°C/14h dark 18°C). 6 plants were grown in boxes containing 500 mL of media. The media was renewed weekly. After 5 weeks, shoots were harvested, dried at 88 °C for 20 h, and digested with concentrated nitric acid. Digested samples were diluted with 18 MΩ water and analyzed by using inductively coupled plasma (ICP)-MS (NexION 2000; PerkinElmer).

### Natural soil microbiome

For the root microbiome analysis on natural soil we used sandy loam (pH 6.7, 1.4 % organic carbon) from the University of Warwick Crop Centre (Wellesbourne, UK). Seeds were sown in 1.5 in × 1.5 in pots and stratified for two days at 4°C. Seeds from the different lines were sown using a randomised scheme and plants were grown under controlled environment conditions (12 h light/12 h dark, 22°C) for four weeks. DNA was extracted from rhizosphere samples (root and adhering soil) using the Griffith method (Griffith et al., 2000). DNA was normalised to 5 ng/μl before PCR amplification. For each sample, 15 ng of DNA was used to amplify part of the 16S rRNA gene using primer pairs 515f and 806r (Caporaso et al., 2011). The primer set was modified at the 5′ end with Illumina Nextera Index Kit v2 adapters. PCR reactions were performed in a reaction volume of 25 μl, containing Q5® Hot Start High-Fidelity 2X Master Mix (New England Biolabs) and 0.5 μM of each primer. Thermocycling consisted of an initial denaturation at 98°C for 30 s followed by 25 cycles of 98°C for 10 s, 50°C for 15 s and 72°C for 20 s with a final extension at 72°C for 5 min. Following PCR, DNA amplicons were purified using Agencourt AMPure XP beads (Beckman Coulter, USA) according to the manufacturer’s instructions. The adapted amplicons were then modified by attaching indices and Illumina sequencing adapters using the Nextera XT Index Kit v2 by PCR as described in the manufacturer’s protocol, enabling simultaneous sequencing of multiple samples. Following the index PCR, the DNA amplicons were purified and normalised using the SequalPrep™ Normalization Plate (96) Kit (Invitrogen) and then quantitatively assessed using a Qubit 2.0 Fluorometer (Life Technologies, USA). The final concentration of the pooled library was 4 nM. The library was sequenced using the MiSeq Reagent Kit v3 600-cycle (Illumina).

Following sequencing, Trimmomatic v0.35 was used to remove low-quality bases from the sequence ends(Bolger et al., 2014). The following steps were then performed using USEARCH and UPARSE software(Edgar, 2010, 2013). Paired-ends reads were assembled by aligning the forward and reverse reads, trimming primers and quality filtering (-fastq_maxee 0.5). Unique sequences were sorted by abundance and singletons were discarded from the dataset. Sequences were clustered to OTUs at 97% minimum identity threshold using -cluster_otus, which includes chimera filtering. This was followed by further chimera filtering using the gold database(Edgar et al., 2011). Sequences were clustered to OTUs at 97% minimum identity threshold. Taxonomy was assigned using Quantitative Insights into Microbial Ecology (QIIME 1.8) and the Greengenes reference database(Caporaso et al., 2010; McDonald et al., 2012). Mitochondrial and chloroplast sequences were removed from the dataset. A total 896,362 sequence reads were assigned to bacteria and used in subsequent analyses with an average of 12,805 reads per sample.

The Phyloseq package (version 1.6.0) in the R software environment was applied for data interpretation and graphical representation including nonmetric multidimensional scaling (NMDS)(McMurdie and Holmes, 2013).

### SynCom microbiome analysis

The effect of host genotype on plant-associated bacterial community structure was tested using recolonization experiments with a gnotobiotic system based on calcined clay inoculated with a SynCom of 206 At-RSPHERE root isolates complemented with 30 soil abundant isolates (Bai et al., 2015). In brief, 100 g of dry sterile calcined clay in Magenta boxes was mixed with 70 ml ½ MS medium containing carrier solution (10 mM MgSO_4_) and/or the bacterial SynCom, resulting in about 2.75*106 cells per g calcined clay.

One-week-old seedlings were transferred from germination plates to the Magenta boxes at 4 seedlings per box and further incubated for 7 weeks. For each plant genotype, three independently prepared SynComs were inoculated in three technical replicates. At harvest, rosette, roots and clay of unplanted boxes were harvested and DNA was isolated (including the input SynComs) using the FastDNA kit for Soil after milling harvested materials in a 2 ml lysing matrix E tube (MP Biomedicals). To assess the bacterial community structure, a 16S rRNA gene-based analysis was employed. The variable regions V5-V7 of bacterial 16S rRNA genes were amplified by PCR. In a second PCR, Illumina adapters and barcodes were added to products of the first PCR. All samples were gelpurified and pooled in equimolar amounts and the final amplicon libraries were twice purified using Agencourt AMPure XP beads (Beckman Coulter) and sequenced on the Illumina MiSeq platform using the MiSeq Reagent kit v3 (2x 300 bp pair-end reads and 12 bp barcode).

Forward and reverse reads were joined, demultiplexed and subjected to quality control using a combination of QIIME and USEARCH pipelines (Caporaso et al., 2010; Edgar, 2010). High quality sequences were clustered to 99% sequence identity together with reference sequences of root and soil isolates assembled in the SynCom using the UPARSE algorithm (Edgar, 2013). OTUs corresponding to one or more reference 16S rRNA gene sequence(s) were selected and an OTU table was generated with read frequencies. Beta-diversity measures (Bray-Curtis) were calculated and canonical analysis of principle coordinates was computed on the distance matrix with the capscale function implemented in the vegan R package. Significance of the constrained variable ‘genotype’ was calculated by a permutation-based ANOVA test using 1000 permutation. Additionally, relative abundances of the 10 most abundant families (based on the average family abundance across all samples) were calculated and visualized as stacked barplots using ggplot2. Analysis of variance with Student-Newman-Keuls, as post-hoc test as implemented in the agricolae R package, was used to assess significant differences in relative abundance.

## Acknowledgements

We thank Gary Grant for help with plant husbandry; Niko Geldner, Barbara Kobei and Isabelle Carre for seed stocks and reagents; Alex Jones for advice on phosphoproteomic analysis; Liliana M. Costa for discussions and comments on the manuscript. Supported by JSPS KAKENHI grant (17H03782) to T.K., ERC AdG ROOTMICROBIOTA, CEPLAS and Max Planck Society to P.S.L. and BBSRC grants (BB/N023927/1 and BB/L027739/1) to D.S and (BB/L025892/1) to G.B and (BB/L003023/1, BB/N005279/1, BB/N00194X/1 and BB/P02601X/1) to J.G-M.

## Author contributions

D.S., G.B. and J.G-M. defined strategy, supervised and procured funding. J.D. and G.R. conducted genetic, molecular and histological analyses. J.D. and S.H. performed the natural soil root microbiome analysis. J.D and S.S. performed the SynCom root and leaf microbiome analysis. J.D. and T.K. designed and constructed the ERK1-GFP and BSK1-GFP reporter lines. J.D. and A.F. designed and constructed vectors for recombinant protein expression. H.G.D. performed the electron microscopy analysis. P.F. performed IC-PMS analysis. J.D. and J.G-M. wrote the paper with input from all authors.

## Data deposition

The mass spectrometry proteomics data have been deposited to the ProteomeXchange Consortium via the PRIDE partner repository with the dataset identifier PXD013861 and 10.6019/PXD013861. Sequencing reads have been deposited at the NCBI SRA Archive (accession no. PRJNA543397).

## Supplementary figures

**Supplementary figure 1. *ERK1* and *RBK1* expression, T-DNA lines and *erk1-3* complementation.**

A Expression of *ERK1* (AT5G65530).

B Expression of *RBK1* (AT5G10520).

C Gene models of *ERK1* and *RBK1*. T-DNA lines used in this study are indicated with triangles.

D Quantification of PI penetration into the stele quantified as number of endodermal cells from the first fully expanded cell (n = 15).

**Supplementary figure 2. Transmission electron microscopy analysis of wild type and *erk1-3* roots.**

A, C Cellular organisation and CS cell wall deposition in root endodermis of wild type plants.

B, D Cellular organisation and CS cell wall deposition in root endodermis of *erk1-3* plants.

**Supplementary figure 3. Quantitative phosphoproteomic analysis in erk1-3 roots identified TIC and TOL6 as downstream targets.**

A Protein model of TIC and TOL6. Serine sites differentially phosphorylated between

*erk1-3* and Col-0 are indicated.

B Normalised abundance of differentially phosphorylated TIC peptide.

C Normalised abundance of differentially phosphorylated TOL6 peptide.

D Gene ontology analysis of proteins differentially phosphorylated in erk1-3 roots (> 1.5-fold change, p-value 0.001).

**Supplementary figure 4. Apoplastic barrier integrity in TOL and circadian clock mutants.**

A and B Quantification of PI penetration into the stele quantified as number of endodermal cells from the first fully expanded cell (n = 10).

**Supplementary figure 5. Lignin deposition in different casparian strip mutants and different mutant combinations.**

**Supplementary figure 6. Quantification of PI penetration into the stele of endodermal cells of casparian strip mutants and different mutant combinations.**

**Supplementary figure 7. Quantification of suberinisation of endodermal cells in different casparian strip mutants.**

**Supplementary figure 8. Distribution of lignin and CASP1-GFP in roots of casparian strip mutants upon exposure to CIF2 peptides.**

A Lignin deposition in root endodermis after exposure to CIF2 peptides.

B CASP1-GFP deposition in root endodermis after exposure to CIF2 peptides.

**Supplementary figure 9. Distribution of suberin in roots of casparian strip mutants upon exposure to CIF2 peptides.**

**Supplementary figure 10. Inonome analysis and germination under hyperosmotic salt stress conditions.**

A Principal component analysis based on the inomic concentration of 16 elements in shoots.

B – D Germination rate of seed under different salt concentrations (n=6, 200 seed per sample).

**Supplementary figure 11. Effect of excess iron in vegetative growth of different casparian strip mutants.**

**Supplementary figure 12. Bacterial communities of inoculated with SynCom cultured communities.**

A Canonical analysis of principal coordinates (based on Bray–Curtis distances) showing different leaf-associated communities of SynComs (n = 10).

A Canonical analysis of principal coordinates (based on Bray–Curtis distances) showing different root-associated communities of SynComs (n = 10).

C The relative abundances of the 10 most abundant families were statistically compared within one organ (root / leaf) using analysis of variance with Student-Newman-Keuls as post-hoc test. Families without a letter annotation do not differ significantly (p > 0.05).

