## Supplementary figures and images for "Two Receptor-Like Kinases Required For Arabidopsis Endodermal Root Organisation Shape The Rhizosphere Microbiome"

### Supplemental Figures

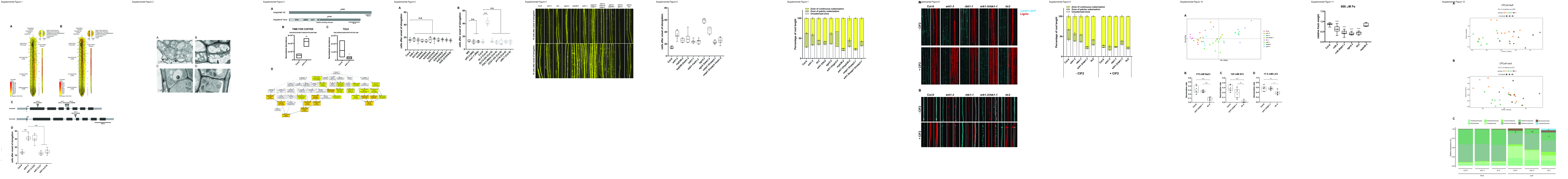
